# Minimum flow decomposition guided by saturating subflows

**DOI:** 10.64898/2025.12.11.693570

**Authors:** Ke Chen, Abhishek Talesara, Sanchal Thakkar, Mingfu Shao

## Abstract

The minimum flow decomposition problem abstracts a set of key tasks in bioinformatics, including metagenome and transcriptome assembly. These tasks, collectively known as multi-assembly, aim to reconstruct multiple genomic sequences from reads obtained from mixed samples. The reads are first organized into a directed graph (e.g., overlap graph, splice graph), where each edge has an integer weight representing the number of supporting reads. By viewing the graph as a flow network, the underlying sequences and their abundances can be extracted through decomposition into a minimum number of weighted paths. Although this problem is NP-hard, prior work has proposed an efficient heuristic that transforms the graph by identifying nontrivial equations in the flow values. However, for graphs with complex structures, many equations cannot be fully resolved by existing mechanisms, leading to suboptimal decompositions. In this study, we revisit the theoretical framework of the flow decomposition problem and extend the equation-resolving mechanisms to jointly model all equations in the graph, enabling safe merge operations that iteratively simplify the graph. Experimental results demonstrate that our new algorithm substantially improves decomposition quality over existing heuristics, achieving near-optimal solutions for complex graphs, while running several orders of magnitude faster than the ILP formulation. Source code of our algorithm is available at https://github.com/Shao-Group/catfish-LP.

## 1 Introduction

Minimum flow decomposition (MFD) is a fundamental problem in graph theory that asks to decompose a directed acyclic graph with a unique source *s*, a unique sink *t*, and a valid flow on its edges into a minimum set of weighted *s*-*t* paths whose combined contributions exactly explain the given flow. MFD abstracts a broad class of computational biology tasks collectively known as multi-assembly. In the general setting, sequencing reads are obtained from an unknown collection of sequences with unknown counts and are aggregated into an assembly graph; the goal is then to reconstruct those sequences and estimate their relative abundances. A prominent example is reference-based transcriptome assembly [24, 11, 23, 22, 16, 18, 15, 19, 29, 25]. For each gene, mapped RNA-seq reads are combined into a splice graph whose edge weights represent the number of reads supporting each splice junction. Decomposing this graph into *s*-*t* paths recovers the expressed RNA isoforms from the gene. Similarly, the decomposed paths can represent contigs or strain haplotypes from a microbial community in metagenomic assembly [21]; or viral strains inferred from infected hosts in the study of virus quasispecies [1].

Despite being a clean and idealized theoretical model for the above multi-assembly tasks, MFD is computationally challenging. It is known to be strongly NP-hard [26] and hard to approximate within some fixed constant factor [12]. To compute optimal solutions, Kloster et al. [14] proposed a fixed-parameter tractable (FPT) algorithm whose running time grows exponentially in the number of paths in the solution. More recently, a series of studies introduced integer linear programming (ILP) formulations for MFD [5, 9], which can handle larger instance sizes than the FPT approach and are versatile enough to incorporate extensions such as inexact flows [27, 4], safety and subpath constraints [7, 13, 28, 3], as well as graphs with cycles [6]. However, despite improvements in modeling and steady progress in commercial solvers such as Gurobi [10], the intrinsic NP-hardness of ILP limits their scalability on large practical datasets.

A complementary line of research focuses on heuristics that aim to produce practically acceptable decompositions efficiently. Among these, the greedy-width algorithm [26], which iteratively extracts the path with maximum possible flow, is widely used and often outperforms methods with stronger theoretical guarantees [12], even though it can be exponentially worse than optimal in the worst case [2]. This disconnection between theoretical guarantees and empirical performance suggests a major gap in our understanding of the structure of MFD and the behavior of its algorithms. A major step toward bridging this gap was made in catfish [20], where an efficient heuristic was introduced based on analyzing the linear algebraic structures of optimal decompositions. By identifying and resolving linear equations among edge flows, catfish substantially improves decomposition quality on both simulated and real transcriptomic graphs. The graph instances introduced in [20] have since become a standard benchmark, and even the most advanced ILP formulations still struggle to solve all of them within a reasonable amount of time.

While efficient and generally superior to greedy-width, catfish remains vulnerable to suboptimal solutions due to the presence of superficial equations—linear relations between the edge flows that are not supported by any minimum decomposition. Distinguishing good equations from superficial ones often requires global structural information that local flow values alone cannot capture. Our work addresses this limitation through two key insights. First, many structural dependencies among edge flows can be tested using an efficient linear programming (LP) formulation that checks whether the subflows saturating each edge are compatible with a candidate set of linear equations. Note that, unlike ILP, LP can be solved in polynomial time, so this filtering step incurs only modest computational overhead. Second, while the primary goal is to determine whether the LP system is feasible, the feasible solution returned by the solver often reveals additional opportunities to further simplify a complex graph, making it possible to tackle equations that were previously unresolvable. Building on these observations, we propose catfish-LP (Algorithm 2), an LP system that models the saturating subflows required for optimal decomposition. This single LP formulation is highly versatile, enabling more effective filtering of superficial equations, safer edge-merging operations, and a principled LP-guided greedy extraction strategy. Through extensive experiments, we show that catfish-LP produces significantly higher-quality decompositions than previous heuristic methods, while remaining orders of magnitude faster than state-of-the-art ILP formulations.

## 2 Preliminaries

We first formally define the MFD problem. Let *G* = (*V, E*) be a directed acyclic graph with a unique source vertex *s*, a unique sink vertex *t*, and an integral flow *f* : *E* → ℤ. Let *P* be the set of all *s*-*t* paths in *G*. A subset of paths *P* ⊆ *P* together with an integral weight function *w* : *P* → ℤ^+^ is called a decomposition of the flow network (*G, f*) if the weighted paths in *P* fully account for the flow in *G*, namely, 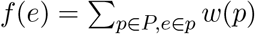 for all *e* ∈ *E*. The minimum flow decomposition problem asks for a decomposition (*P, w*) that minimizes the number of paths |*P* | in it. See an example in Figure 1. Next, we recall a few key concepts from [20] that are relevant to catfish-LP.

**Figure 1:**
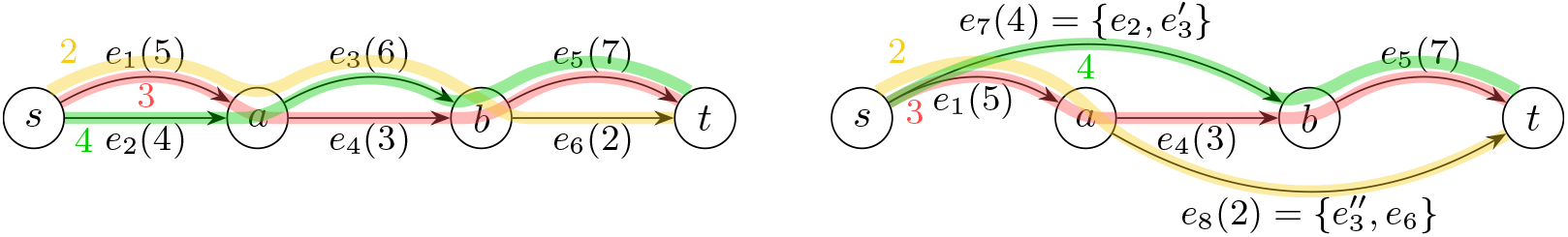
An example of reducing the optimality gap by resolving an equation. The flow value for each edge is shown in parentheses. Left: Observe that Δ = 6 − 4 + 2 = 4. An optimal (minimum) decomposition (*P* ^∗^, *w*^∗^) has three paths: 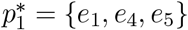 with weight 3, 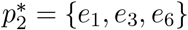 with weight 2, and 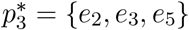 with weight 4. So the optimality gap is 1. The matrix *P* ^∗^ has a nontrivial null vector (0, 1, −1, 0, 0, 1)^*T*^ . It corresponds to an equation of flow values *f* (*e*_2_) + *f* (*e*_6_) = *f* (*e*_3_). Right: The equation can be resolved by splitting *e*_3_ into 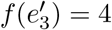 and 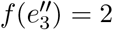, merging *e*_2_ with 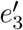 to get *e*_7_, and merging 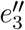 with *e*_6_ to get *e*_8_. In the resulting graph, Δ = 5 − 4 + 2 = 3, so the optimality gap has been reduced from 1 to 0. Applying a greedy algorithm on this graph can successfully produce the optimal decomposition *P* ^∗^.

### 2.1 Null space of a set of paths

Assigning arbitrary indices *E* = {*e*_1_, *e*_2_, …, *e*_|*E*|_} and *P*= {*p*_1_, *p*_2_, …, *p*_|P|_} allows us to write *P*as a binary matrix of size |P| × |*E*|, where P[*i, j*] = 1 if *p*_*i*_ contains *e*_*j*_ and 0 otherwise. Consequently, any subset of paths *P* ⊆ *P* can be viewed as a binary matrix of size |*P* | × |*E*| by keeping only the rows of *P*corresponding to the paths in *P* . We slightly abuse notation by using *P*(and *P*) to denote both the set of paths and the corresponding matrix; the intended meaning should be clear from context. A column vector *q* with |*E*| entries is called a null vector of a matrix *P* if *P* · *q* = **0**. The set of all such vectors forms the null space *N* (*P*) = {*q* | *P* · *q* = **0**}. For each vertex *v* ∈ *V* − {*s, t*}, its incidence vector *q*_*v*_ ∈ {0, ±1}^|*E*|×1^ is defined by setting *q*_*v*_[*i*] = 1 if *e*_*i*_ enters *v, q*_*v*_[*j*] = −1 if *e*_*j*_ leaves *v*, and *q*_*v*_[*k*] = 0 for all remaining entries. It is straightforward to verify that every *q*_*v*_ is a null vector for any matrix *P* . We denote by *N*_0_ the linear space spanned by the set {*q*_*v*_ | *v* ∈ *V* − {*s, t*}} and call vectors in *N*_0_ trivial null vectors.

### 2.2 Optimality gap

Let (*P* ^∗^, *w*^∗^) be a minimum decomposition. An upper bound |*P* ^∗^| ≤ Δ = |*E*| − |*V* | + 2 is known from [26]. In [20], the authors define the optimality gap to be Δ − |*P* ^∗^| and show that it is positive if and only if there exist null vectors outside of *N*_0_.

#### Proposition 1

(Proposition 4 in [20])

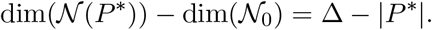

See Figure 1 (Left) for an example.

Since the upper bound Δ applies to any decomposition algorithm that produces independent paths, including the greedy algorithm that repeatedly selects and removes the *s*-*t* path with the largest flow value, reducing the optimality gap can potentially improve the performance of the greedy algorithm. Our goal is to apply transformations to the input flow network that reduce Δ without altering the (unknown) optimal solution. An example is shown in Figure 1 (Right).

### 2.3 Equations of flow values

Both *f* and *w* can be represented as integral row vectors, and an equivalent formulation of (*P, w*) being a valid decomposition of (*G, f*) is simply *f* = *w*·*P* . Let (*P* ^∗^, *w*^∗^) be a minimum decomposition. For any null vector *q* ∈ N (*P* ^∗^), we have *f* · *q* = (*w*^∗^ · *P* ^∗^) · *q* = *w*^∗^ · (*P* ^∗^ · *q*) = *w*^∗^ · **0** = 0. Thus, each null vector of *P* ^∗^ induces a linear equation among the edge flow values, providing a way to reason about the structure of an optimal decomposition without explicitly knowing it.

Restricting attention to null vectors with entries in {0, ±1} yields equations involving two disjoint sets of edges, corresponding to coefficients +1 and −1. For example, in Figure 1 (Left), the nontrivial null vector (0, 1, −1, 0, 0, 1)^*T*^ gives rise to the equation *f* (*e*_2_) + *f* (*e*_6_) = *f* (*e*_3_). Such an equation has a clear structural interpretation: there exists an optimal decomposition in which every path using an edge on the left-hand side must also use some edge on the right-hand side. In this example, the equation implies that all flow through *e*_2_ must eventually pass through *e*_3_, resolving the ambiguity at node *a* in Figure 1 and allowing *e*_2_ and *e*_3_ to be safely merged. Equations of this type provide strong, structure-based guidance toward the underlying (unknown) optimal decomposition. The catfish algorithm [20] builds directly on this observation by iteratively identifying and resolving such equations through local graph transformations. To handle equations involving non-adjacent edges, catfish introduces a mechanism known as the *reverse operation*, which transforms the graph structure while preserving the set of feasible flow decompositions. Reverse operations operate on closed subgraphs and flip their internal orientations to make distant edges adjacent, while maintaining a one-to-one correspondence between decompositions of the original and transformed graphs.

For completeness, we outline the original catfish algorithm in Algorithm 1, which will serve as a conceptual foundation for the methods developed in subsequent sections.

#### Algorithm 1

Catfish

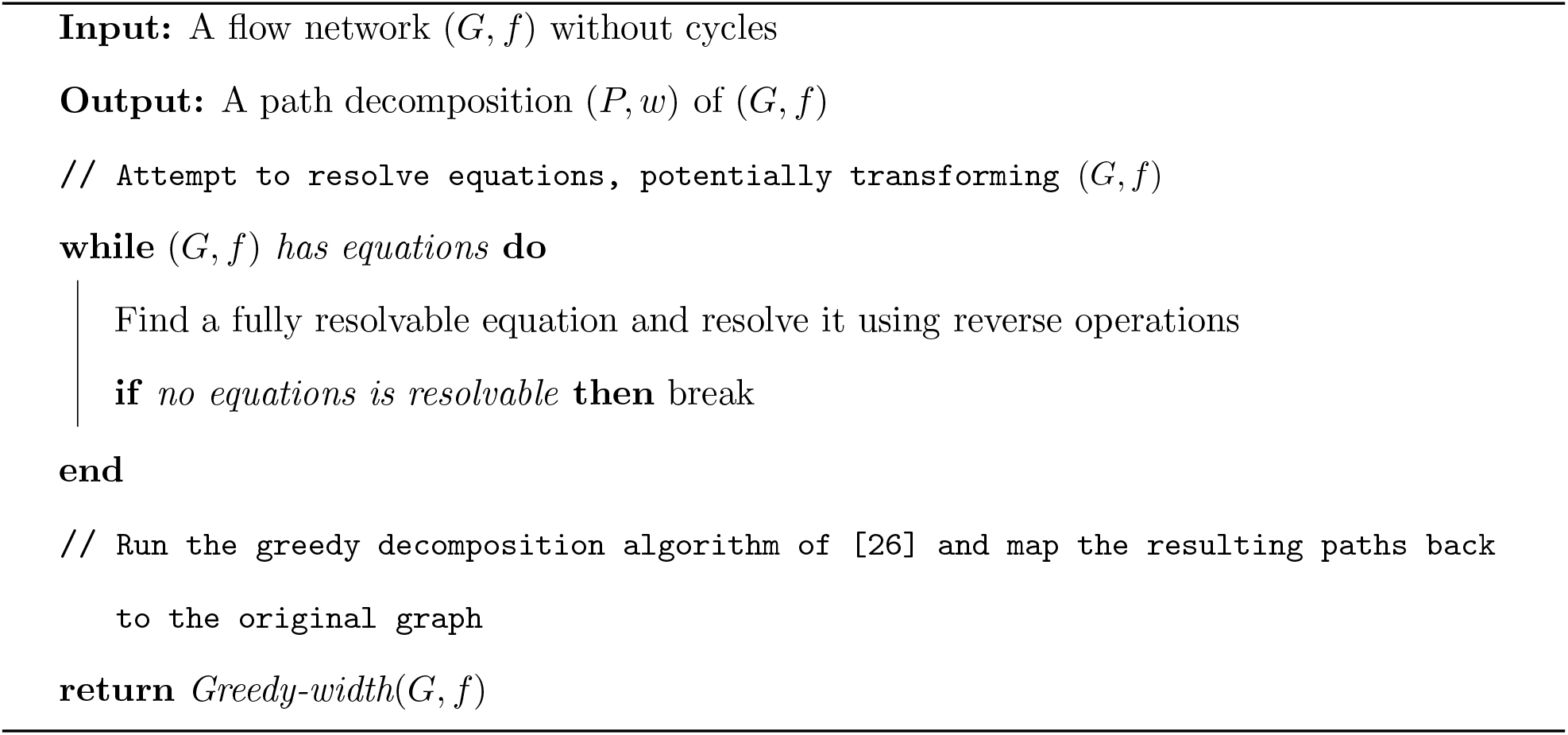

## 3 Limitations of catfish

The while-loop in catfish reduces the optimality gap by iteratively resolving equations on edge flows. While efficient and empirically effective on simpler graphs, catfish exhibits three fundamental limitations that restrict its performance on more complex instances.

### 3.1 Superficial equations hurt decomposition

The equation-resolving mechanisms in catfish provide an effective way to exploit good equations, but they rely critically on the assumption that the equations being resolved correspond to nontrivial null vectors of an optimal decomposition. The difficulty is that this correspondence is only one-directional. Although every nontrivial null vector induces a linear equation on edge flow values, the converse does not hold: a flow graph may satisfy linear equations that do *not* correspond to any null vector of a minimum decomposition. We call an equation *superficial* if it does not arise from any nontrivial null vector, and *good* otherwise.

For example, in Figure 1 (Left), the equation *f* (*e*_1_) + *f* (*e*_6_) = *f* (*e*_5_) corresponds to the vector *q* = (1, 0, 0, 0, −1, 1)^*T*^, which is not in N (*P* ^∗^) (e.g., 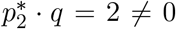). Resolving such equations is therefore not justified by the structure of any minimum decomposition.

In [20], it was shown that under certain uniformity assumptions, most equations encountered during the execution of catfish are good and informative, particularly when the graph structure is not overly convoluted, which explains its strong empirical performance on simpler instances. In more complex graphs, however, superficial equations are more likely to be selected, and resolving them can distort the graph structure in ways that are incompatible with any minimum decomposition, leading to suboptimal results (see Figure 2 for an example). This exposes a key missing component in the original framework: a principled mechanism for validating whether an equation is consistent with all feasible decompositions.

**Figure 2:**
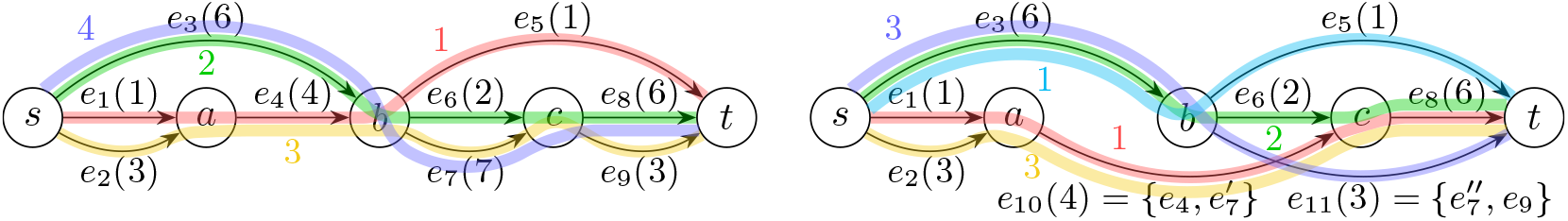
An example where resolving a superficial equation leads to suboptimal decomposition. Left: Original graph with four paths in a minimum decomposition: 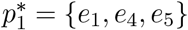 with weight 1, 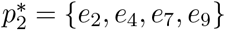 with weight 3, 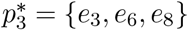 with weight 2, and 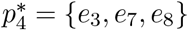with weight 4. Right: After resolving the superficial equation *f* (*e*_4_) + *f* (*e*_9_) = *f* (*e*_7_) by splitting *e*_7_(7) into 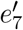 (4) and 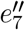 (3) and merging with *e*_4_ and *e*_9_ respectively, the new graph requires five paths in a minimum decomposition.

### 3.2 Fully resolving an equation is difficult

While the equation-resolving mechanisms of catfish come with strong theoretical guarantees, namely, that optimal decompositions are preserved, they also limit the algorithm’s effectiveness in practice. In particular, in complex graphs, closed subgraphs on which reverse operations can act are often rare, making it difficult or impossible to fully resolve many equations.

When an equation cannot be fully resolved, catfish discards all information provided by that equation. In the extreme case where no equation is fully resolvable, the algorithm falls back entirely to greedy decomposition. This behavior trades safety for effectiveness: although partial resolution may be safe in some cases, catfish lacks a systematic mechanism for determining which aspects of an unresolvable equation can be applied without violating optimality, and therefore opts to ignore them altogether. This limitation suggests the need for a framework that can exploit partial information from good equations in a principled manner.

### 3.3 *P*remature fallback to greedy decomposition

A related limitation is that catfish may revert to greedy decomposition even when the graph still contains equations with substantial structural information. Empirically, greedy decomposition often performs poorly on complex instances, producing solutions with sizes close to the theoretical upper bound Δ and leaving a large optimality gap.

In such situations, a more effective strategy would be to perform an *informed greedy* decomposition—one that remains consistent with the constraints implied by the unresolved equations—rather than abandoning equation-based guidance entirely. The absence of such a mechanism further limits the performance of catfish on challenging graphs.

These observations motivate the need for a more robust framework that can validate equations, exploit partial information, and guide decomposition beyond purely combinatorial rules.

## 4 Method

Rather than introducing ad hoc heuristic fixes for each limitation of catfish in isolation, we address them in a unified manner by augmenting its equation-resolving framework with a lightweight yet expressive linear programming (L*P*) formulation, named catfish-L*P*. The key idea is to use L*P* not as an exact solver for MFD, but as a structural oracle that validates equations, extracts consistent subflow information, and guides safe graph transformations.

### 4.1 The base model

Let *m* = |*E*| be the number of edges in *G*. The base model introduces *m*^2^ continuous variables *x*_*e*_(*a*), indexed by ordered pairs of edges *e, a* ∈ *E*. Intuitively, *x*_*e*_(·) is intended to represent a valid *s*-*t*-subflow in *G* that saturates edge *e*; equivalently, it encodes the collective flow of all paths in a decomposition that traverse *e*. We enforce this interpretation through the following constraints:

#### Constraint 1

*x*_*e*_(*e*) = *f* (*e*) for all *e* ∈ *E*: the subflow associated with *x*_*e*_(·) saturates edge *e*.

#### Constraint 2

L 0 ≤ *x*_*e*_(*a*) ≤ *f* (*a*) for all *e, a* ∈ *E*: each edge in each subflow carries a valid amount of flow.

#### Constraint 3

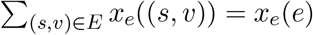 the subflow defined by *x*_*e*_(·) must all pass through the edge *e*.

#### Constraint 4

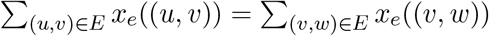 for all *e* ∈ *E* and all *v* ∈ *V* − {*s, t*}: each subflow satisfies flow conservation at every intermediate vertex.

Note that the above constraints, particular Constraints 3 and 4, already imply

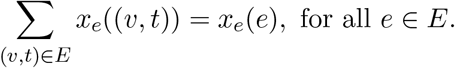

In other words, the subflow saturating edge *e* must eventually all reach the sink *t*. We therefore do not include these constraints explicitly.

We also do not impose constraints of the form 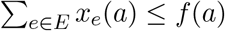, which would require the total flow through edge *a* across all subflows to remain within its capacity. Such constraints are invalid because the subflows are allowed to overlap. For example, if a path in *P* ^∗^ saturates both edges *e*_1_ and *e*_2_, then the corresponding subflows defined by 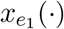 and 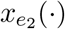 coincide along the entire path. In particular,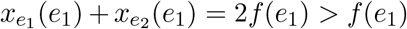, showing that the aggregate contribution of multiple subflows may exceed the edge capacity.

At the same time, subflows must not be entirely independent. To encode this dependency, we further impose the following symmetry constraint:

#### Constraint 5

*x*_*e*_(*a*) = *x*_*a*_(*e*) for all edges *e, a* ∈ *E*: both variables represent the total amount of flow that passes through both edges *e* and *a*.

The above constraints collectively model necessary conditions satisfied by any optimal decomposition (*P* ^∗^, *w*^∗^). For each edge *e*, aggregating all weighted paths in *P* ^∗^ that traverse *e* yields a solution for *x*_*e*_(·) that satisfy all above constraints. In particular, for any two edges *e, a* ∈ *E*,

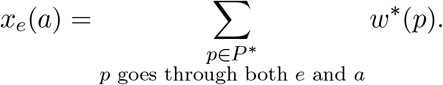

Thus, *x*_*e*_(*a*) represents the amount of flow on edge *e* that must pass through edge *a*, for all pairs of edges. Equivalently, the solution encodes pairwise co-occurrence information between edges under the optimal decomposition.

However, these base constraints capture only necessary conditions and may therefore admit solutions that do not correspond to an optimal decomposition. (From this point of view, the above base model serves as a relaxation of the optimal decomposition.) To further restrict the feasible region, we introduce the following equation constraints, which steer the L*P* toward the structure of the optimal decomposition.

### 4.2 Equation constraints

We consider only equations on edge flow values with coefficients in {0, ±1}. Such an equation can be specified by two subsets *E*_*L*_, *E*_*R*_ ⊆ *E*: writing 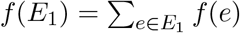for any *E*_1_ ⊆ *E*, the form *f* (*E*_*L*_) = *f* (*E*_*R*_) (see Figures 1 and 2 for examples). Since linear combinations of equations are also equations, we restrict attention to indivisible equations—those that admit no strict subsets 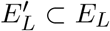and 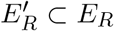 with 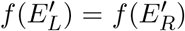—because they are smaller in size and already encode all relevant information. Note that being indivisible implies that *E*_*L*_ and *E*_*R*_ are disjoint. If an indivisible equation *f* (*E*_*L*_) = *f* (*E*_*R*_) indeed arises from a nontrivial null vector of some minimum decomposition *P* ^∗^, then it captures the structural dependency that any path in *P* ^∗^ using an edge in *E*_*L*_ must also use some edges in *E*_*R*_, and vice versa. Moreover, the subflow saturating an edge in *E*_*L*_ should not use any other edges of *E*_*L*_ (same for *E*_*R*_). To model these relationships, we add the following 2*m* constraints for an equation:

#### Constraint 6

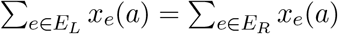 for all *a* ∈ *E*: for each edge, the total subflow through edges in *E*_*L*_ must equal that through edges in *E*_*R*_.

#### Constraint 7

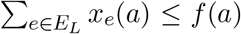 for all *a* ∈ *E*: the combined subflow contributed by edges in *E*_*L*_ (and therefore *E*_*R*_) must not exceed the total flow on each edge.

This L*P* formulation checks a necessary condition for the equation *f* (*E*_*L*_) = *f* (*E*_*R*_) to correspond to a nontrivial null vector of a minimum decomposition *P* ^∗^: if the equation is good with respect to *P* ^∗^, then the induced subflows saturating each edge (obtained by aggregating all paths in *P* ^∗^ that use the edge) satisfy all the above constraints, and hence the L*P* model is feasible. Taking the contrapositive, an infeasible L*P* instance certifies that the equation cannot arise from a nontrivial null vector of any minimum decomposition, and thus the equation should be discarded.

Better yet, this formulation also allows us to test whether a set of equations is compatible (since all good equations arising from nontrivial null vectors of *P* ^∗^ must be mutually compatible). Starting from the base model, we can add equation constraints sequentially and check feasibility at each step. If the LP becomes infeasible after adding a particular equation, that equation (and its associated constraints) is removed. This procedure further filters out superficial equations, preventing structurally invalid resolutions from distorting the graph and thereby improving the quality of the final decomposition.

Our LP formulation can effectively leverage the graph structures, flow constraints, and equation constraints all together to filter out superficial equations. As a concrete example, consider the left graph in Figure 2. Observe that Constraint 6 applied to the equation *f* (*e*_1_) = *f* (*e*_5_) requires all flow on *e*_1_ to eventually pass through *e*_5_. Since the path *e*_1_ → *e*_4_ → *e*_5_ is the only path capable of satisfying this requirement, we can conclude that it must appear in the final decomposition. The equations *f* (*e*_2_) = *f* (*e*_9_) and *f* (*e*_3_) = *f* (*e*_8_) are both compatible with this requirement while the superficial equation *f* (*e*_4_) + *f* (*e*_9_) = *f* (*e*_7_) is not. Indeed, one unit of flow on *e*_4_ is forced to continue through *e*_5_, and hence cannot all pass through *e*_7_. Consequently, this equation is rejected, preventing the suboptimal decomposition shown in Figure 2 (Right).

It is worth noting that the LP model could be strengthened to check for more conservative conditions, for example, by adapting the ILP formulation of [5]. However, our experiments indicate that the proposed LP model is orders of magnitude more efficient, and its filtering quality is already sufficient for practical use.

### 4.3 Transform *G* with a feasible LP

The LP-based verification can be directly integrated into Algorithm 1. The primary remaining bottleneck is that not all good equations can be successfully resolved with existing mechanisms, especially when the graph structure is complex. A feasible LP model, after filtering out superficial equations, provides valuable structural information that can be used to simplify the graph. When no equation can be fully resolved, we apply the following two heuristics:

1. Recall that a feasible LP solution provides, for each edge *e* ∈ *E*, a valid subflow *x*_*e*_(·) that saturates *e*. If, in this subflow, an edge *e*^′^ satisfies *x*_*e*_(*e*^′^) = *f* (*e*), then there exists a valid decomposition (consistent with all remaining equations) in which every unit of flow through *e* also uses *e*^′^. Moreover, if *e* and *e*^′^ are adjacent, they can be conveniently merged while preserving the decomposition. However, this merge may be only supported by some feasible decompositions discovered by the LP and is not guaranteed to be correct for an optimal one. To err on the safe side, we again use LP to determine whether *e* and *e*^′^ must be merged in all decompositions consistent with the current equation set. This can be done by adding a temporary constraint *x*_*e*_(*e*^′^) *< f* (*e*) − *ϵ* and test for feasibility. If the LP becomes infeasible, the merge is deemed safe, and we contract *e* and *e*^′^ to simplify the graph. Such merges can have meaningful downstream effects, for example, enabling the resolution of equations that are otherwise unresolvable.
2. If no further safe simplifications are possible, Algorithm 1 exits the while-loop and proceeds to greedy decomposition. Empirically, we observe that greedy decomposition almost always yields inferior solutions when unresolvable equations remain. In such cases, the feasible LP solution itself may lead to a better outcome. We examine the LP solution to identify any saturating subflow *x*_*e*_(·) that is actually a simple path through *e*. Among all such paths, we extract the one carrying the maximum flow *f* (*e*), remove it from the graph, and add it to the final decomposition. Unlike the previous heuristic, this operation is not guaranteed to be safe as the extracted path may not appear in all compatible decompositions. Instead, it serves as an “informed greedy” step: selecting the heaviest path that is consistent with all remaining equations, rather than the heaviest path in the graph overall.

In summary, catfish-LP employs a structure-based LP model that, within a unified framework, addresses the three limitations of catfish described in Section 3. The method orchestrates three components, all derived from the same underlying LP formulation. First, the LP model is used to distinguish good equations from superficial ones by enforcing necessary conditions for compatibility with a minimum decomposition. Second, saturating subflows are extracted from feasible LP solutions, which encode partial yet reliable information implied by good equations and enable safe graph simplifications even when full resolution is not achievable. Finally, the LP solutions guide an informed greedy extraction strategy, deferring the error-prone greedy-width heuristic by prioritizing paths that remain consistent with all good equations. The complete algorithm is outlined in Algorithm 2.

#### Algorithm 2

Catfish-LP

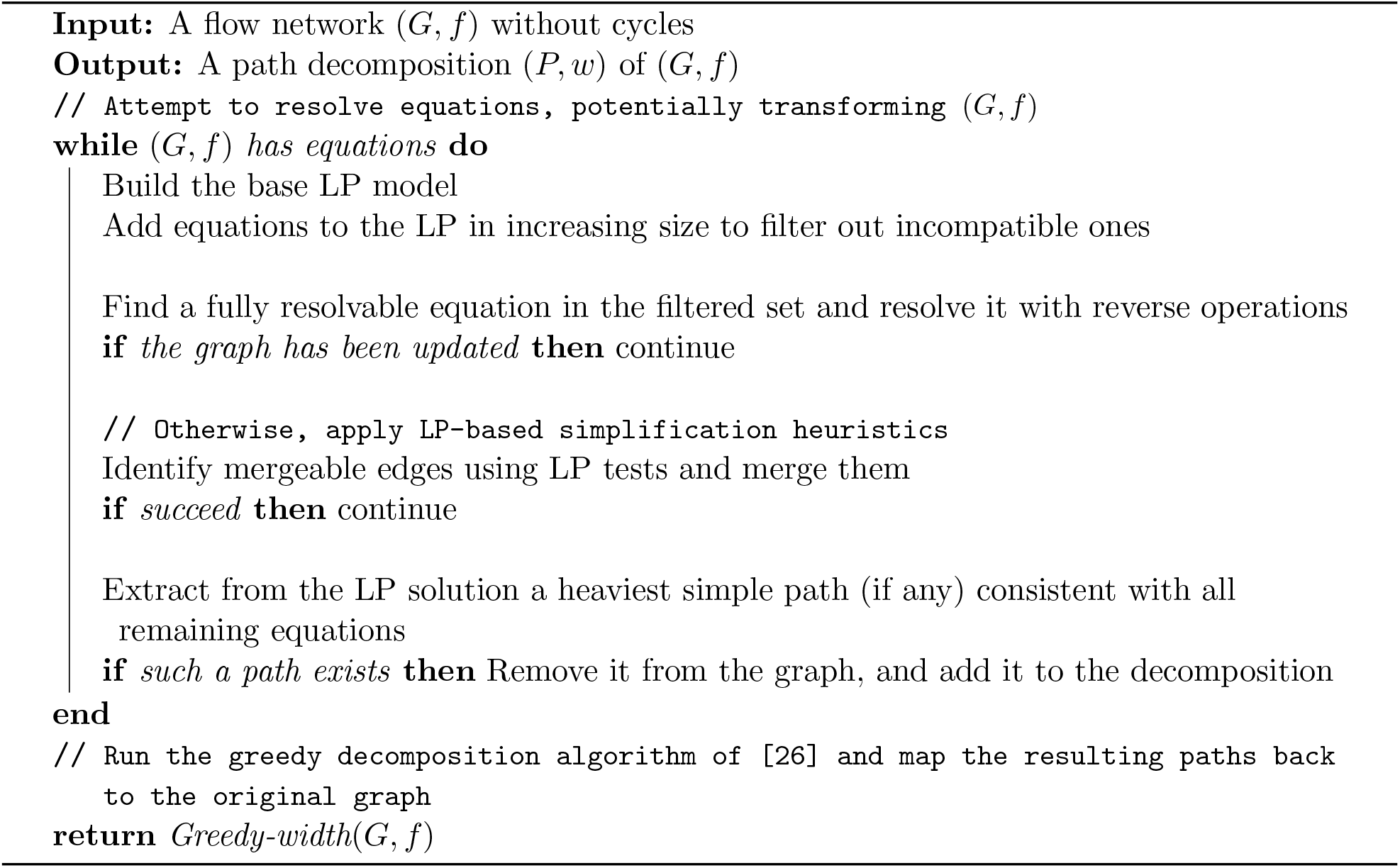

## 5 Experimental results

We evaluate four algorithms for the MFD problem: the greedy-width and catfish algorithms as implemented in [20], the optimized ILP formulation from [9], and our enhanced catfish-LP. All experiments are conducted on an Ubuntu server equipped with an Intel(R) Xeon(R) Gold 6148 CPU @ 2.40GHz and 566 GB RAM. Both ILP and catfish-LP rely on the same Gurobi installation; ILP uses the Python API whereas catfish-LP uses the C++ interface. Although ILP performs model construction in Python and catfish-LP in C++, the impact of the implementation language on overall performance is negligible, since the Gurobi solver dominates the runtime in both cases. Each method is limited to at most five cores per instance. Greedy-width and catfish are entirely single-threaded. ILP and catfish-LP are also single-threaded outside the solve, but their Gurobi calls utilize multiple cores. The five-core limit was chosen to match the computational resources of our experimental setup. Because we report total CPU time recorded using time(1) -v rather than wall-clock time, variations in parallelism do not affect the measured performance. For each instance, ILP requires a valid initial decomposition to enable its safety optimizations, for which we supply the greedy-width solution. Since the running time of greedy-width is negligible compared to ILP, its execution time is not included in ILP’s total time. A wall-clock time limit of 30 minutes is imposed for each instance.

### 5.1 Results on perfect splice graphs

We first evaluate all methods on a now-standard benchmark introduced by Shao and Kingsford [20]. It includes four datasets of perfect splice graphs. The first dataset is derived from human RNA-seq samples quantified using Salmon [17]. For each gene in each sample, perfect splice graphs are constructed by superimposing all expressed transcripts with their estimated abundances. We refer to this dataset as Salmon.

The remaining three datasets are generated via simulation using the Flux-Simulator [8] on three well-annotated species: human, mouse, and zebrafish. We refer to these datasets by the corresponding species name.

Table 1 summarizes the decomposition quality of each method on the four datasets. Because the input graphs are constructed from ground-truth paths, we use the total number of such paths, |*P* ^†^|, as a baseline for evaluating all algorithms. Note that |*P* ^†^| does not necessarily correspond to the minimum possible decomposition; indeed, both Catfish-LP and ILP frequently produce even smaller solutions. However, the ground-truth paths are biologically meaningful: in transcript assembly, pursuing a strictly minimum decomposition is somewhat arbitrary, whereas recovering the true underlying isoforms is a more relevant objective (though difficult to assess on real data where ground truth is unavailable). For this reason, we also report the precision and recall of each algorithm in Table 2. Precision is defined as the fraction of reconstructed paths that exactly match a ground-truth path, while recall is defined as the fraction of ground-truth paths that are successfully recovered. A reconstructed path is considered correct only if both its node sequence and its weight exactly match those of a ground-truth path.

**Table 1:** Decomposition results on perfect splice graphs. The column |*P* ^†^| shows the total number of ground-truth paths (RNA isoforms) in each dataset. For the ILP column, the number in parentheses is the number of ground-truth paths in instances that ILP timed out.

| Dataset | Graphs | $\vert P^\dagger\vert $ | $\vert P\vert - \vert P^\dagger\vert $ | | | |
| --- | --- | --- | --- | --- | --- | --- |
|  |  |  | greedy-width | catfish | catfish-LP | ILP |
| Salmon | 13,300,893 | 32,889,145 | 1,330,137 | 86,355 | 15,093 | -11,072 (43,476) |
| human | 1,169,083 | 2,139,559 | 13,252 | 491 | -142 | -763 (169) |
| mouse | 1,316,058 | 2,142,500 | 11,410 | -1 | -817 | -1,332 (233) |
| zebrafish | 1,549,373 | 2,140,200 | 2,928 | -387 | -386 | -508 (16) |

**Table 2:**
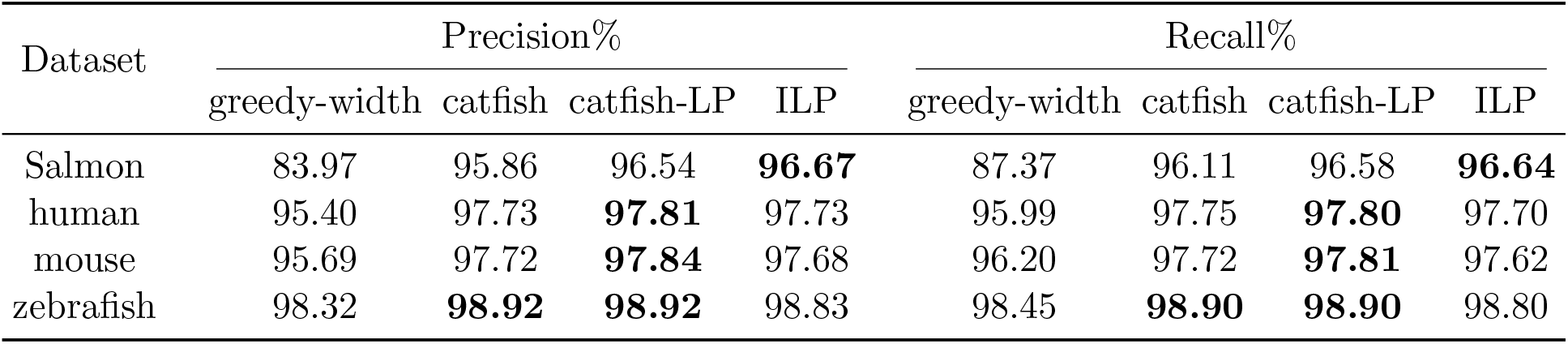
Precision and recall on perfect splice graphs. The best results are highlighted for each dataset.

Observe that catfish-LP outperforms both greedy-width and catfish in producing smaller de-compositions. Although ILP appears to obtain even smaller decompositions, it timed out on some of the most complex instances (see also Table 3), which artificially improves its reported numbers. At the same time, catfish-LP consistently achieves the highest precision and recall on all datasets except Salmon, suggesting that the saturating subflows it relies on successfully capture meaningful structural information within splice graphs.

**Table 3:** Running time on perfect splice graphs. The ILP timeout column shows the number of instances/total number of ground-truth paths in those instances, for which ILP failed to produce a decomposition within the 30 minutes time limit.

| Dataset | CPU time (seconds) |  |  |  | ILP timeout |
| --- | --- | --- | --- | --- | --- |
| | greedy-width | catfish | catfish-LP | ILP | Graphs/$\vert P^\dagger\vert $ |
| Salmon | 2,713 | 2,945 | 158,862 | 24,182,561 | 2,106/43,476 |
| human | 174 | 159 | 7,772 | 179,445 | 12/169 |
| mouse | 178 | 152 | 8,820 | 216,275 | 15/233 |
| zebrafish | 195 | 151 | 9,752 | 13,421 | 1/16 |

Table 3 reports the CPU time of all methods on the four datasets. As expected, catfish-LP is slower than catfish due to the additional LP-based filtering, which is the key contributor to its improved decomposition quality. Nonetheless, because LP is polynomial-time solvable, catfish-LP remains orders of magnitude faster than ILP. Among the four methods, only ILP exceeds the 30-minute time limit on certain instances, suggesting that its true time to completion would, in many cases, be even larger.

### 5.2 Results on simulated graphs

As reported in [20], and as reflected in the second and third columns of Table 1, most perfect splice graphs in the above datasets are relatively simple: the vast majority admit ground-truth decompositions of at most ten paths. To better assess algorithmic performance on more challenging instances, we generated additional synthetic graphs with 20 or 30 vertices, between 10 and 30 ground-truth paths, and maximum path lengths ranging from 10 to 23. Increasing the number and length of ground-truth paths on a fixed vertex set naturally creates more entangled graph structures, making accurate decomposition substantially harder. This dataset contains 1, 440 simulated graphs, comprising 20 graphs for each of the 72 configurations (details in Appendix A), and it allows us to systematically evaluate algorithm performance across a range of graph complexities and difficulty levels.

Figure 3 summarizes both decomposition quality and running time on this simulated dataset. Observe that catfish-LP consistently produces smaller decompositions and recovers more ground-truth paths than all other heuristic methods. ILP, by contrast, fails to finish within the time limit on a large fraction of the more complex graphs—precisely the instances that also challenge the heuristic methods—making its results less directly comparable in this experiment. This is also evident from its runtime profile: nearly 90% of ILP’s total reported CPU time is spent in timeout instances. On the remaining, easier graphs where ILP does terminate, its total runtime is 524,656 seconds, whereas catfish-LP requires less than 3% of that time, or 15,553 seconds, to achieve comparable decomposition quality. See Figures 4 and 5 for more details.

**Figure 3:**
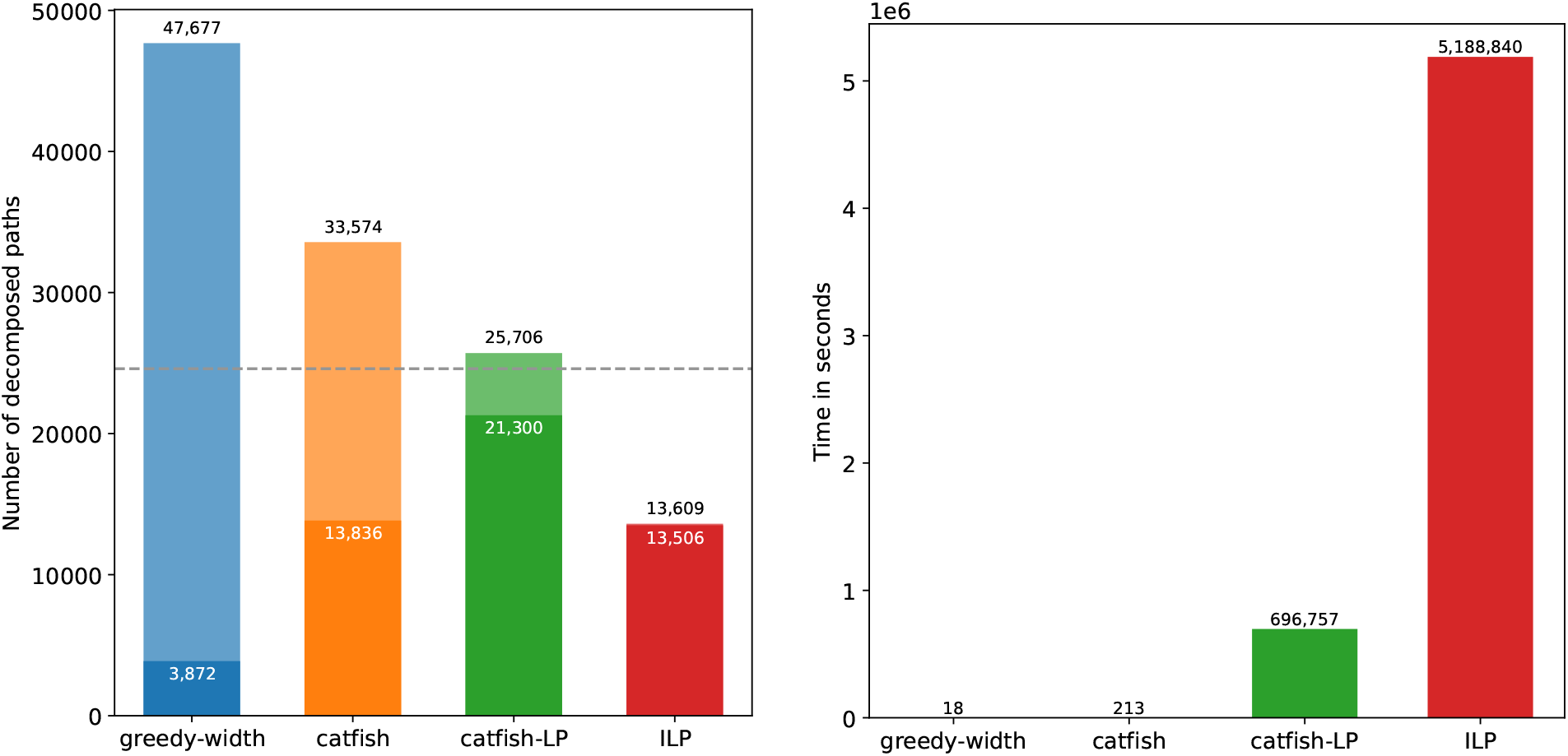
Decomposition quality and runtime on the full simulated dataset. In the left subfigure, the dashed horizontal line marks the total number of ground-truth paths across all instances (|*P* ^†^| = 24, 600). For each method, the full bar represents the total number of paths produced, while the shaded portion denotes the number of decomposed paths that exactly match a ground-truth path. The ILP bar is heavily skewed because ILP failed to decompose 498 graph instances (35%) within the time limit.

**Figure 4:**
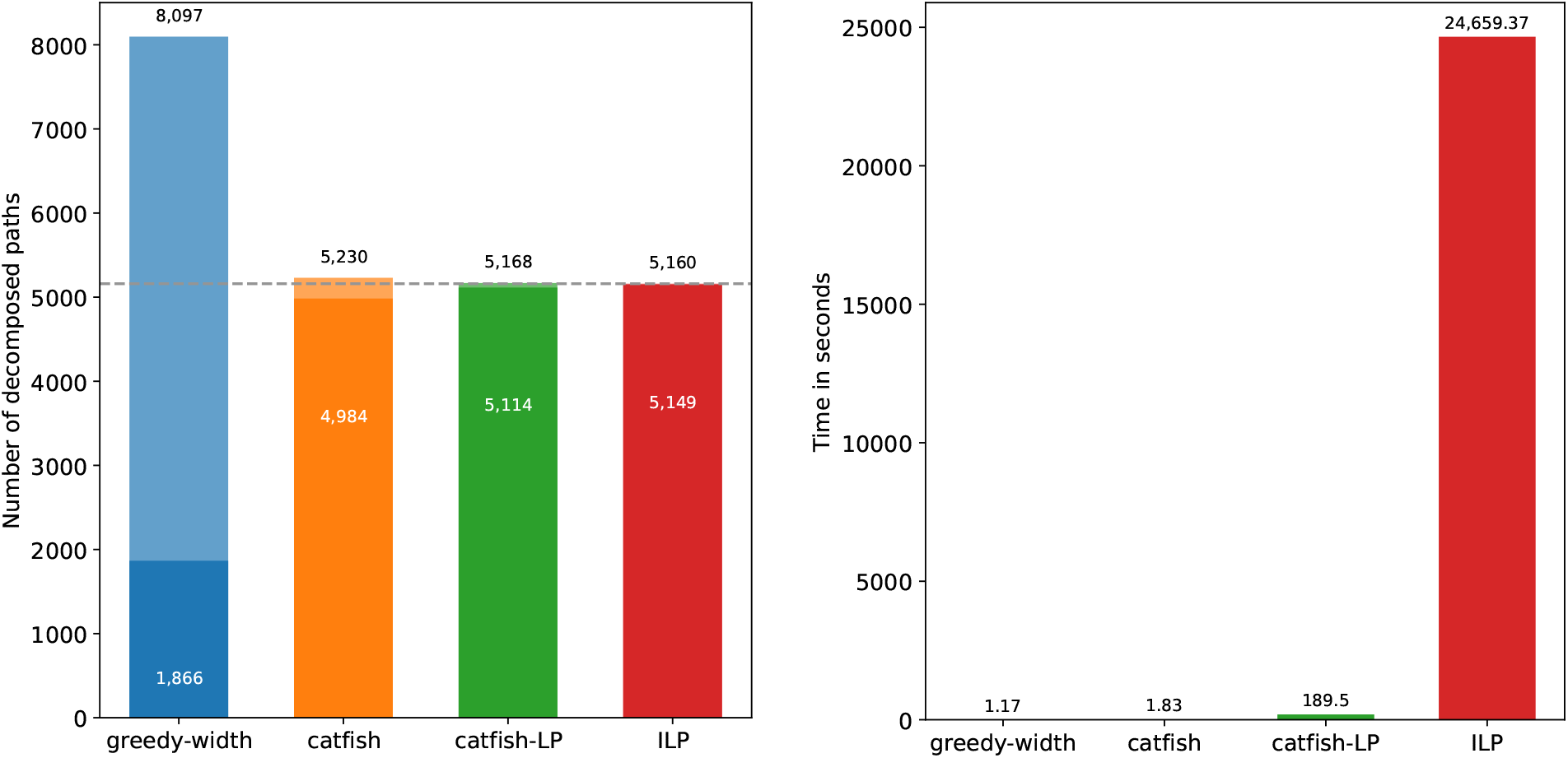
Decomposition quality and runtime on the subset of 22 configurations where ILP finished all 20 instances within time limit. In the left subfigure, the dashed horizontal line marks the total number of ground-truth paths across all instances (|P_†_| = 5, 160).

**Figure 5:**
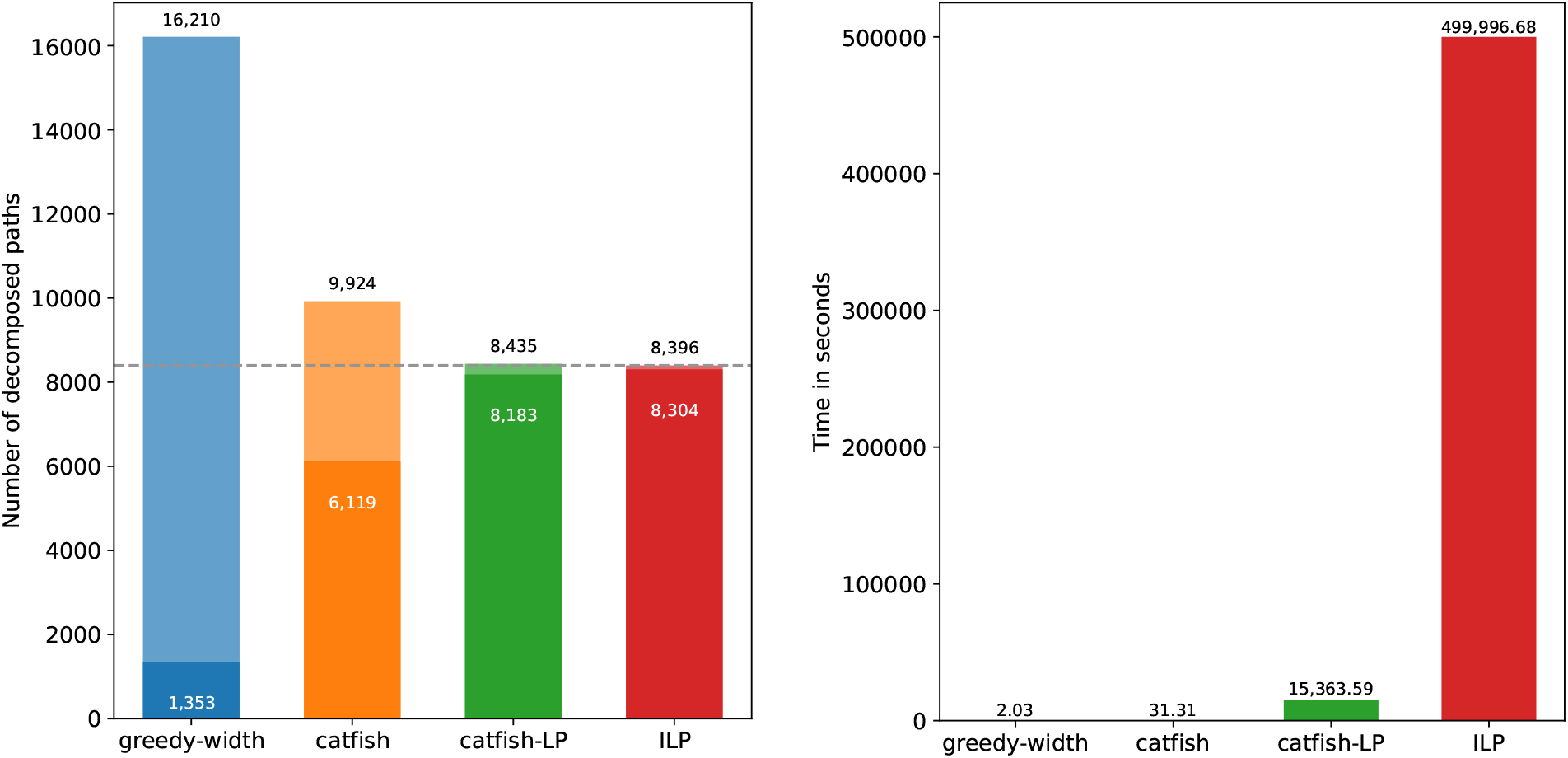
Decomposition quality and runtime on harder configurations where ILP finished some instances within time limit. There are 502 graphs and the number of ground-truth paths is |*P* ^†^| = 8, 396.

For a fairer comparison, we report in Figure 4 both decomposition quality and running time on the subset of 22 configurations in which ILP successfully completed all 20 instances within the time limit. This subset corresponds to relatively easier instances, on which the original catfish already achieves near-perfect decomposition performance. Even in this setting, catfish-LP further improves upon catfish by reducing the total number of reconstructed paths while also increasing the number of correctly recovered ground-truth paths. Its decomposition quality is within 1% of ILP, while it completes all instances in 3 minutes compared to 6.8 hours for ILP.

Next, we consider a more challenging subset of graphs corresponding to configurations in which ILP completed only a subset of instances within the time limit. This dataset contains 502 graphs from 43 configurations, with a total of 8, 396 ground-truth paths. The results are shown in Figure 5. On this harder dataset, catfish exhibits a noticeable drop in decomposition quality, whereas catfish-LP achieves substantial improvements and remains on par with ILP whiling using only 3% of its runtime.

For the remaining 498 most challenging graphs, on which ILP timed out on every instance, we report the decomposition quality and runtime of greedy-width, catfish, and catfish-LP in Figure 6. On these instances, catfish struggles to produce high-quality decompositions for two reasons. First, the number of superficial equations increases substantially, interfering with the graph simplification heuristics. Second, the increasingly tangled graph structure makes it difficult to fully resolve the good equations even if they are identified. As a result, the performance of catfish degrades towards that of greedy-width. In contrast, catfish-LP is able to capture substantially more structural information to guide the decomposition process. Consequently, it achieves a threefold improvement in the number of recovered ground-truth paths while reducing the total number of reconstructed paths by roughly one third.

**Figure 6:**
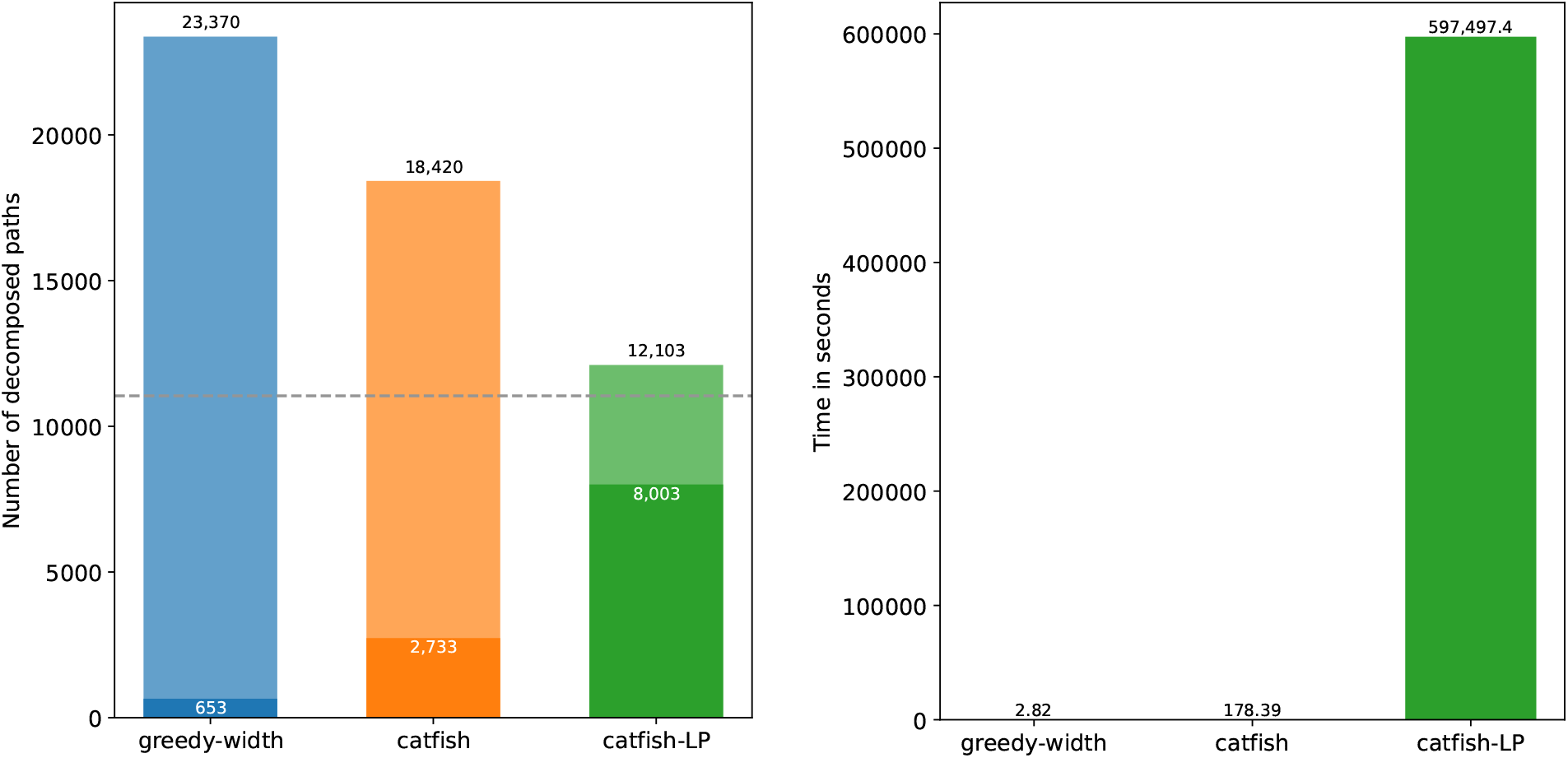
Decomposition quality and runtime on the hardest instances where ILP timed out. There are 498 graphs and the number of ground-truth paths is |*P* ^†^| = 11, 044.

## 6 Summary and discussion

This work introduced catfish-LP, a new algorithm for the minimum flow decomposition problem that augments the state-of-the-art heuristic framework, catfish, with a lightweight yet highly informative use of linear programming. By leveraging LP both as a filter for identifying structurally consistent equations and as a guide for graph simplification via saturating subflows, catfish-LP addresses a central challenge in the theoretically motivated equation-resolving-based strategy: determining which equations are good for an unknown optimal decomposition and how to exploit them when direct resolution fails. Unlike ILP-based approaches, which offer exact solutions but scale poorly, our LP-based design remains computationally efficient while capturing much of the structural insight needed for high-quality decompositions.

Across a broad collection of datasets, including splice graphs derived from real RNA-seq data and a controlled suite of simulated graphs with diverse levels of complexity, catfish-LP consistently produces decompositions that are nearly as small as those obtained by ILP while achieving substantially higher recall, outperforming classical heuristics such as greedy-width and the original catfish. These improvements validate the central premise of the method: feasible LP solutions expose structurally meaningful saturating subflows that can validate good equations, guide safe graph transformations, and enable informed greedy extractions in situations where purely combinatorial heuristics frequently stuck or make suboptimal choices.

Our experiments further show that catfish-LP provides a compelling balance between accuracy and scalability. Although slower than the purely combinatorial catfish algorithm due to the necessary LP invocations, it remains orders of magnitude faster than ILP and scales effectively to large and entangled graphs where ILP routinely hits time limits. Moreover, the LP formulation introduced here provide a flexible foundation that can be adapted to other MFD variants with meaningful bioinformatics applications.

Overall, catfish-LP demonstrates that incorporating LP into the combinatorial structure of MFD yields both practical benefits and conceptual insights. Practically, it produces smaller and more accurate decompositions within realistic computational budgets. Conceptually, it highlights how LP-feasible subflows can encode structural information about the decomposition space, suggesting new directions for hybrid optimization-combinatorial approaches for MFD and related flow-based problems. Future directions include exploring strengthened LP formulations, integrating domain-specific constraints, and developing probabilistic interpretations of LP-guided decompositions for transcriptome assembly and beyond.

## 7 Acknowledgments

This work is supported by the US National Science Foundation (2145171 to M.S.) and by the US National Institutes of Health (R01HG011065 to M.S.).

## A Configurations of simulated graphs

The 72 configurations used in Section 5.2 are listed in the following table.

| Configuration | Number of vertices | Number of paths | Maximum path length |
| --- | --- | --- | --- |
| 1 | 20 | 10 | 10 |
| 2 | 20 | 10 | 11 |
| 3 | 20 | 10 | 12 |
| 4 | 20 | 10 | 13 |
| 5 | 20 | 10 | 14 |
| 6 | 20 | 10 | 15 |
| 7 | 20 | 10 | 16 |
| 8 | 20 | 12 | 10 |
| 9 | 20 | 12 | 11 |
| 10 | 20 | 12 | 12 |
| 11 | 20 | 12 | 13 |
| 12 | 20 | 12 | 14 |
| 13 | 20 | 12 | 15 |
| 14 | 20 | 12 | 16 |
| 15 | 20 | 14 | 10 |
| 16 | 20 | 14 | 11 |
| 17 | 20 | 14 | 12 |
| 18 | 20 | 14 | 13 |
| 19 | 20 | 14 | 14 |
| 20 | 20 | 14 | 15 |
| 21 | 20 | 14 | 16 |
| 22 | 20 | 16 | 10 |
| 23 | 20 | 16 | 11 |
| 24 | 20 | 16 | 12 |
| 25 | 20 | 16 | 13 |
| 26 | 20 | 16 | 14 |
| 27 | 20 | 16 | 15 |
| 28 | 20 | 16 | 16 |
| 29 | 20 | 18 | 10 |
| 30 | 20 | 18 | 11 |
| 31 | 20 | 18 | 12 |
| 32 | 20 | 18 | 13 |
| 33 | 20 | 18 | 14 |
| 34 | 20 | 18 | 15 |
| 35 | 20 | 18 | 16 |
| 36 | 20 | 20 | 10 |
| 37 | 20 | 20 | 11 |
| 38 | 20 | 20 | 12 |
| 39 | 20 | 20 | 13 |
| 40 | 20 | 20 | 14 |
| 41 | 20 | 20 | 15 |
| 42 | 20 | 20 | 16 |
| 43 | 30 | 10 | 15 |
| 44 | 30 | 10 | 17 |
| 45 | 30 | 10 | 19 |
| 46 | 30 | 10 | 21 |
| 47 | 30 | 10 | 23 |
| 48 | 30 | 14 | 15 |
| 49 | 30 | 14 | 17 |
| 50 | 30 | 14 | 19 |
| 51 | 30 | 14 | 21 |
| 52 | 30 | 14 | 23 |
| 53 | 30 | 18 | 15 |
| 54 | 30 | 18 | 17 |
| 55 | 30 | 18 | 19 |
| 56 | 30 | 18 | 21 |
| 57 | 30 | 18 | 23 |
| 58 | 30 | 22 | 15 |
| 59 | 30 | 22 | 17 |
| 60 | 30 | 22 | 19 |
| 61 | 30 | 22 | 21 |
| 62 | 30 | 22 | 23 |
| 63 | 30 | 26 | 15 |
| 64 | 30 | 26 | 17 |
| 65 | 30 | 26 | 19 |
| 66 | 30 | 26 | 21 |
| 67 | 30 | 26 | 23 |
| 68 | 30 | 30 | 15 |
| 69 | 30 | 30 | 17 |
| 70 | 30 | 30 | 19 |
| 71 | 30 | 30 | 21 |
| 72 | 30 | 30 | 23 |

## Notes

### Competing Interest Statement

The authors have declared no competing interest.

### Summary of Updates

Revised according to WABI'26 reviewer comments. Added colored decomposition paths in figures 1 and 2; clarified definition of good equations; added more detailed results to Section 5.2; added appendix for configurations of the simulated graphs.

